# ROTACs leverage signaling-incompetent R-spondin for targeted protein degradation

**DOI:** 10.1101/2022.12.08.519573

**Authors:** Rui Sun, Hyeyoon Lee, Christof Niehrs

## Abstract

Proteolysis-targeting chimeras (PROTACs) are an emerging technology for therapeutic intervention but options to target cell surface proteins and receptors remain limited. Here we introduce ROTACs, bispecific WNT- and BMP signaling-disabled R-spondin (RSPO) chimeras, which leverage the specificity of these stem cell growth factors for ZNRF3/RNF43 E3 transmembrane ligases, to target degradation of transmembrane proteins. As proof of concept, we targeted the immune checkpoint protein programmed death ligand 1 (PD-L1), a prominent cancer therapeutic target, with a bispecific RSPO2 chimera, R2PD1. The R2PD1 chimeric protein bound PD-L1 and at picomolar concentration induced its lysosomal degradation. In three melanoma cell lines, R2PD1 induced between 50-90% PD-L1 protein degradation. PD-L1 degradation was strictly dependent on ZNRF3/RNF43. We conclude that signaling-disabled ROTACs represent a novel strategy to target cell surface proteins for degradation.

## INTRODUCTION

Targeted protein degradation (TPD) is a rapidly progressing field that has broadened the scope of therapeutic targets to include historically undruggable proteins and overcome drug resistance (Bekes et al., 2022; Li and Crews, 2022). In contrast to degradation of cytosolic targets, the TPD toolbox to target extracellular proteins is rather limited. To overcome this limitation, LYsosome TArgeting Chimera (LYTAC) were developed that bind extracellular proteins and direct them to lysosomes (Banik et al., 2020). LYTACs rely on transmembrane proteins such as mannose 6-phosphate receptor (Banik et al., 2020) or asialoglycoprotein receptor (Ahn et al., 2021; Zhou et al., 2021) to internalize their cargo. An alternative approach relies on protein targeting to RNF43, a single-pass E3 ubiquitin ligase harboring a ligand-binding ectodomain and an intracellular RING domain (Zebisch et al., 2013). RNF43 is a WNT inhibitor that ubiquitinates the WNT receptors Frizzled and LRP6 to induce their endocytosis and lysosomal degradation (Hao et al., 2012; Koo et al., 2012; Tsukiyama et al., 2015). An antibody-based PROTAC (AbTAC) was developed that employs a bispecific antibody to recruit RNF43 for target protein degradation (Cotton et al., 2021; Marei et al., 2022).

RNF43 and its homologue ZNRF3 are receptors for R-spondins (RSPOs), a family of four secreted stem cell growth factors with crucial implications in multiple biological processes, ranging from development to cancer (Chartier et al., 2016; de Lau et al., 2014; Hao et al., 2016; Kazanskaya et al., 2004; Kazanskaya et al., 2008; Kim et al., 2005; Seshagiri et al., 2012). RSPOs amplify WNT signaling by bridging Leucine-rich repeat containing G protein-coupled receptor 4-6 (LGR4-6) with ZNRF3/RNF43. The ternary complex of LGR, RSPO, and ZNRF3/RNF43 is then internalized and degraded. This effect reduces cell surface E3 ligase activity, which would normally target the WNT receptors for destruction, and enhances WNT signaling (Carmon et al., 2011; de Lau et al., 2014; Glinka et al., 2011; Hao et al., 2016; Hao et al., 2012; Koo et al., 2012). Among the four RSPOs, RSPO2 and RSPO3 are bi-functional ligands, which not only activate WNT signaling but also inhibit BMP signaling (Lee et al., 2020; Sun et al., 2021). Mechanistically, RSPO2/3 bind via their TSP1 domain the BMP receptor type I A (BMPR1A) and bridge the interaction with ZNRF3/RNF43, independent of LGRs (Lee et al., 2020). Once again, the ternary RSPO-BMPR1A-ZNRF3/RNF43 complex internalizes and BMPR1A is degraded in the lysosome, effectively reducing BMP signaling. RSPO binding to ZNRF3/RNF43 (Peng et al., 2013b; Zebisch et al., 2013), LGRs (Peng et al., 2013a; Xu et al., 2013), and BMPR1A (Lee et al., 2020; Sun et al., 2021) is highly modular and relies on the FU1, FU2 and TSP1 domains of RSPOs, respectively. The observation that RSPOs can target different transmembrane receptors (LGRs, BMPR1A) to ZNRF3/RNF43, and that its FU1, FU2, and TSP1 domains neatly separate binding to ZNRF3/RNF43, LGRs, and BMP1RA, respectively, suggests that RSPOs could be engineered as LYTAC for targeted protein degradation via ZNRF3/RNF43 to lysosomes. To explore this possibility, we selected the transmembrane protein PD-L1 as the degradation target. PD-L1 is an immune checkpoint protein expressed in various cancer cells and antigen-presenting cells, while its interaction partner PD-1 is found at the cell surface of T cells (Curiel et al., 2003; Dong et al., 2002; Latchman et al., 2001). Interaction between PD-1 and PD-L1 abrogates T cell receptor-induced signals, preventing antigen-mediated T cell activation, and resulting in poor anti-tumor response (Andrews et al., 2019; Patsoukis et al., 2020). Previous studies showed that lysosomal degradation of PD-L1 can be induced by forced interaction between RNF43 and PD-L1 or by a PD-L1 antibody coupled with a lysosomal targeting motif (Cotton et al., 2021; Zhang et al., 2021).

## RESULTS

### Design and validation of a bi-specific RSPO chimera

We modified RSPO2, which possesses the highest binding affinity to ZNRF3(Park et al., 2018), to generate a bi-specific chimera with simultaneous binding capacity for ZNRF3/RNF43 and PD-L1 (Figure 1A). To prevent unwanted WNT signal activation, we mutated phenylalanine 109 of RSPO2 within the FU2 domain (FA mutation), aiming for a LGR-binding deficient mutant (R2^FA^) (Park et al., 2018). To obtain a PD-L1 binding unit, we cloned the extracellular IgV-like domain of PD-1 (PD1) and generated its high-affinity binding variant (Maute et al., 2015) PD1^HAC^. To obtain the final bi-specific reagent, we fused the RSPO2 FU1/2^FA^ domains with the PD1^HAC^ module via a flexible glycine-serine linker, effectively replacing the TSP1 domain of RSPO2, which is non-essential for ZNRF3/RNF43 binding, resulting in the bi-specific RSPO2 chimera, R2PD1. Replacing the TSP1 domain in RSPO2 also abolishes BMP signaling inhibition (Lee et al., 2020; Sun et al., 2021).

**Figure 1.**
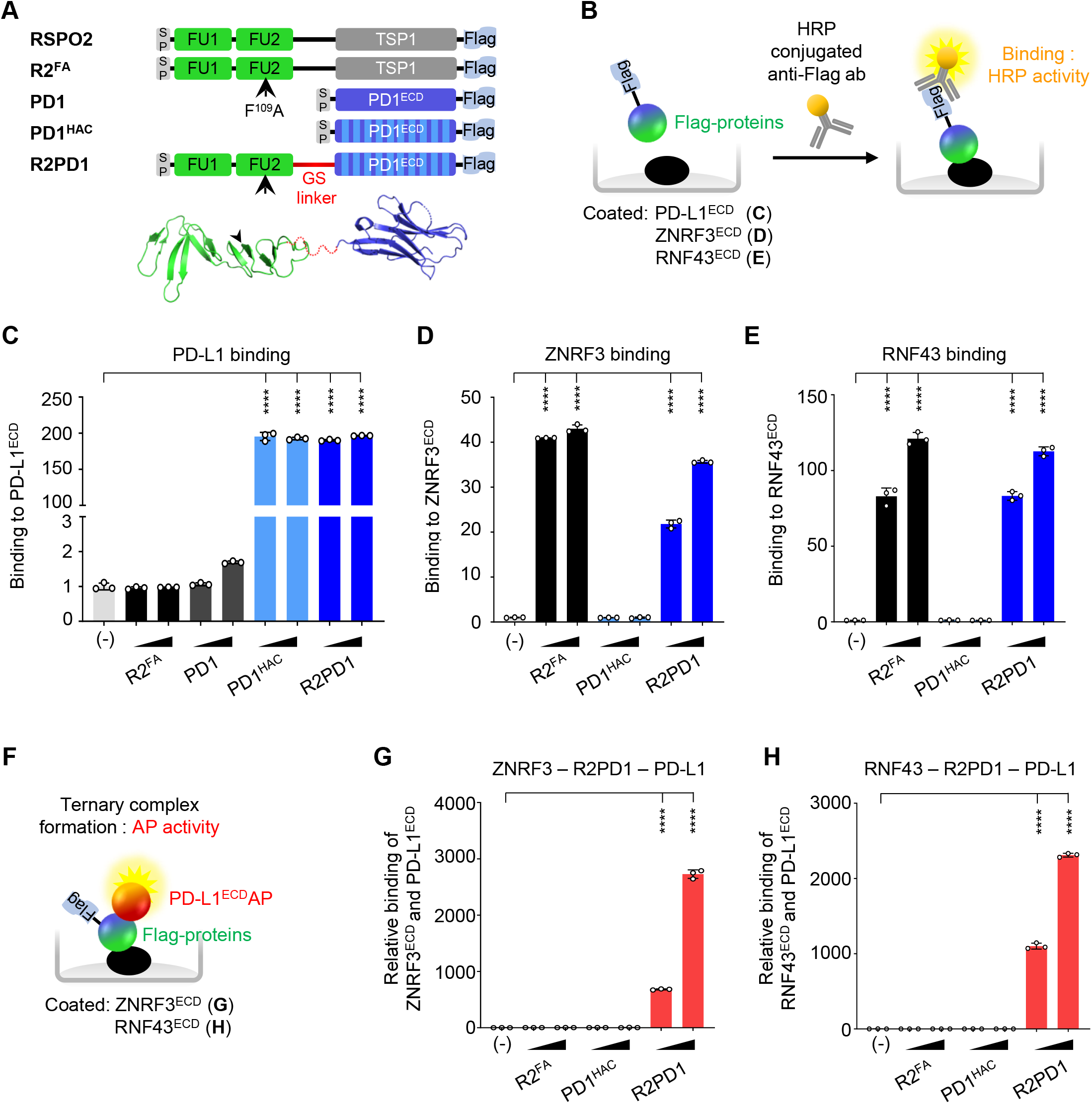
RSPO chimera bridges ZNRF3/RNF43 and PD-L1. **(A)** Top, domain-structure of indicated proteins. Bottom, structural prediction from RSPO2 (PDB 4ufr) and PD-1 HAC (PDB 5ius). R2^FA^, RSPO2 furin domains with a LGR binding deficient F^109^ A mutation. PD1^HAC^, high-affinity consensus (HAC) mutation of PD-1 with enhanced binding to PD-L1. SP, signal peptide; FU, furin domain; GS, glycine-serine linker; TSP1, thrombospondin domain 1. **(B)** Scheme of *in vitro* binding assay in (**C-E**). (**C-E**) *In vitro* binding assay as in (**B**) for direct interaction between RSPO chimera and PD-L1^ECD^ (C), ZNRF3^ECD^ (**D**), RNF43^ECD^ (**E**). n=3 biologically independent samples. Data are displayed as mean ± SD. ****P<0.0001 from two-tailed unpaired t-test. (**F**) Scheme of *in vitro* ternary complex formation assay in (**G** and **H**). (**G, H**) *In vitro* ternary complex formation assay as in (**F**). R2PD1 bridges the interaction between PD-L1^ECD^ and ZNRF3^ECD^ (**G**) and RNF43^ECD^ (**H**). ECD, extracellular domain. n=3 biologically independent samples. Data are displayed as mean ± SD. ****P<0.0001 from two-tailed unpaired t-test. See also **Figure S1**.

Recombinant PD1^HAC^, R2^FA^, and R2PD1 proteins were readily produced and secreted into the medium of transfected HEK293T cells. Expectedly, R2PD1 lost LGR4 binding in a cell surface binding assay (Figures S1A-S1C). To test whether R2PD1 maintains the binding activity towards its interaction partners, we performed *in vitro* binding assays (Figure 1B). Towards immobilized PD-L1, PD1 bound weakly but PD1^HAC^ strongly (Figure 1C), confirming that HAC mutation conveys higher affinity towards PD-L1 (Maute et al., 2015). Importantly, R2PD1 and PD1^HAC^ showed equally strong binding towards immobilized PD-L1 (Figure 1C). Moreover, R2PD1 bound immobilized ZNRF3^ECD^ and RNF43^ECD^ similar to R2^FA^ (Figures 1D and 1E). Thus, R2PD1 retains tight binding towards both its targets, PD-L1 and ZNRF3/RNF43.

The prerequisite of E3 ligase-mediated protein degradation is the formation of an E3 ligase-target complex. To test if R2PD1 bridges the interaction between ZNRF3/RNF43 and PD-L1, we employed *in vitro* binding assays using immobilized ZNRF3^ECD^ and RNF43^ECD^ (Figure 1F). As expected, PD-L1 bound ZNRF3 (Figure 1G) and RNF43 (Figure 1H) only in presence of R2PD1, but not of R2^FA^ or isolated PD1^HAC^ protein. We conclude that RSPO chimera R2PD1 effectively bridges the interaction between PD-L1 and ZNRF3/RNF43.

### RSPO chimera promotes degradation of overexpressed PD-L1

To test whether the RSPO chimera induces PD-L1 degradation, we transfected HEK293T cells with human PD-L1 with or without ZNRF3/RNF43 (Figure 2A). Cotransfection of ZNRF3 or RNF43 had no effect on PD-L1 levels. However, addition of recombinant R2PD1 effectively reduced levels of overexpressed PD-L1, but only when ZNRF3 or RNF43 were cotransfected (Figure 2B). Degradation of PD-L1 reached plateau after incubation with 5.4 nM purified recombinant proteins, with a maximum of 60% total protein reduction detected in an immunoblot (Figure S2A).

**Figure 2.**
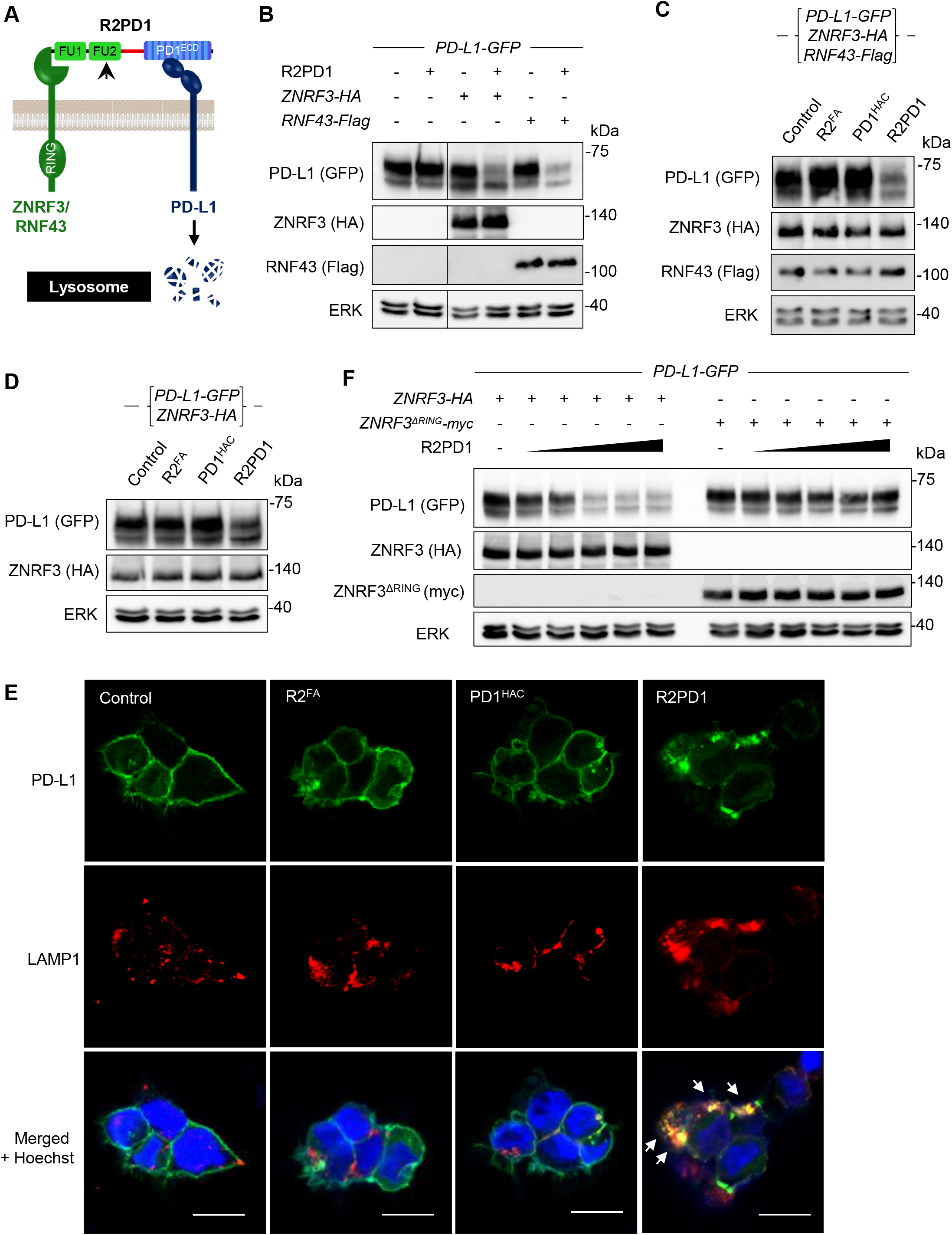
RSPO chimera promotes degradation of overexpressed PD-L1. **(A)** Model of R2PD1 mediated PD-L1 degradation. (**B-D)** Immunoblot analysis of indicated proteins in HEK293T cells. Cells were transfected as indicated and analyzed after 24 h incubation with conditioned medium containing the indicated proteins (**B**-**C**) or 5.4 nM of indicated purified proteins (**D**). Control conditioned medium and mock-purified protein were used as control. **(E)** IF staining of HEK293T cells for indicated proteins. Cells were transfected for 2 days with *PD-L1-GFP* and *ZNRF3-HA*. After additional 5 h incubation with equal amounts of the indicated purified proteins, cells were fixed and stained for GFP and the lysosomal marker LAMP1. Note PD-L1 colocalization with LAMP1 upon treatment with R2PD1 (white arrowheads). Scale bar, 10 μm. **(F)** Immunoblot analysis of indicated proteins in HEK293T cells. Cells were transfected as indicated and analyzed after 24 h incubation with conditioned medium containing the indicated proteins. Control conditioned medium were used as control. See also **Figure S2**.

We hypothesized that PD-L1 degradation requires simultaneous binding to PD-L1 and ZNRF3/RNF43. Indeed, no reduction of PD-L1 levels was detected in cells incubated with equal amounts of either R2^FA^ or PD1^HAC^ (Figures 2C and 2D). This finding was further confirmed by immunofluorescence (IF), where R2PD1 abolished PD-L1 cell surface staining and instead induced colocalization with the lysosomal marker LAMP1 (Figure 2E). No such effect was seen with either R2^FA^ or PD1^HAC^. This result is consistent with previous findings that ZNRF3/RNF43 employ the lysosomal pathway for targeted protein degradation (Koo et al., 2012; Lee et al., 2020). The E3 ligase activity of ZNRF3/RNF43 is crucial for degradation of the WNT receptor Frizzled and BMPR1A (Hao et al., 2012; Lee et al., 2020). To test the importance of E3 ligase activity in concert with PD-L1 degradation, we transfected HEK293T cells with a dominant negative variant of ZNRF3, lacking the intracellular RING domain (ZNRF3^ΔRING^) (Chang et al., 2020; Hao et al., 2012). R2PD1 efficiently reduced total PD-L1 protein level in a dose-dependent manner in cells expressing full-length ZNRF3, while degradation of PD-L1 was abolished upon ZNRF3^ΔRING^ transfection (Figure 2F). Collectively, we conclude that RSPO chimera degrades overexpressed PD-L1 in presence of overexpressed ZNRF3/RNF43 in 293T cells.

### RSPO chimera promotes degradation of endogenous PD-L1 in melanoma cells

To investigate whether RSPO chimera degrades endogenous PD-L1, we turned to cancer cell lines with PD-L1 expression. PD-L1 small interfering RNA (siRNA) knockdown in the human melanoma cell line MEL624 decreased PD-L1 protein levels, confirming both antibody specificity and that the cells express PD-L1 (Figure S3A). Addition of R2PD1 conditioned medium reduced PD-L1 level by ∼ 90% (Figure S3B). PD-L1 reduction plateaued with as little as 0.27 nM R2PD1 protein (Figure 3A). PD-L1 reduction by R2PD1 was readily detected within 6 hours (Figure S3C). Moreover, PD-L1 reduction was only observed with R2PD1, but not R2^FA^ or PD1^HAC^ (Figure 3B), corroborating the requirement for bi-specificity. Flow cytometry analysis confirmed that R2PD1 induced cell surface removal of PD-L1 (Figures 3C and 3D).

**Figure 3.**
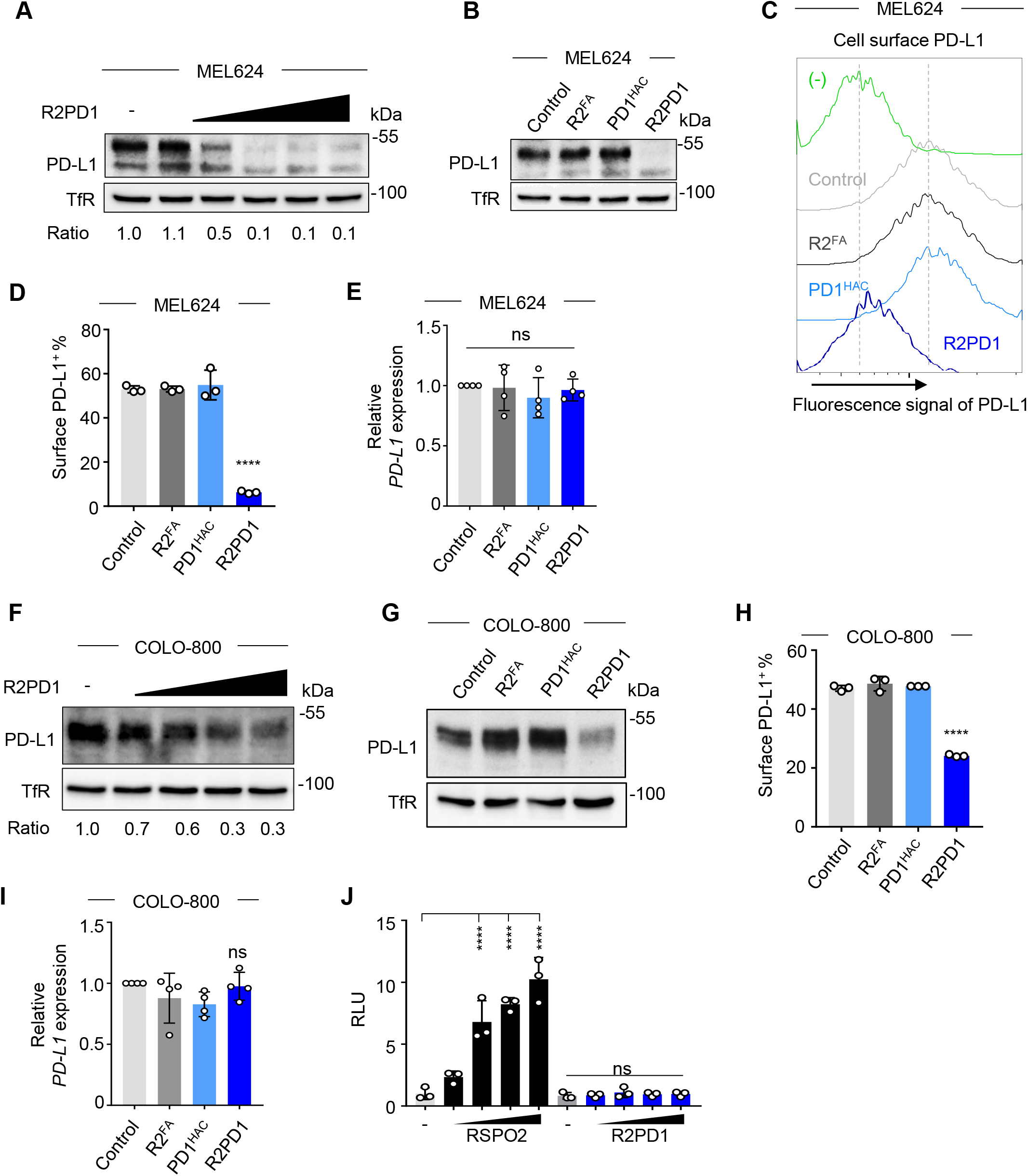
RSPO chimera promotes degradation of endogenous PD-L1 in melanoma cells. (**A-B)** Immunoblot analyses of indicated proteins in MEL624 cells. Cells were analyzed after 24 h incubation with the indicated purified proteins. Mock-purified protein was used as control. The normalized ratio between PD-L1 and TfR is shown below (**A**). **(C)** Flow cytometry analyses of cell surface PD-L1 in MEL624 cells. **(D)** Quantification of (**C**). n=3 biologically independent samples. Data are displayed as mean ± SD. ****P<0.0001 from two-tailed unpaired t-test. **(E)** qRT-PCR analysis of *PD-L1* expression in MEL624 cells. Cells were analyzed after 24 h incubation with the indicated purified proteins. n=4 biologically independent samples. Data are displayed as mean ± SD. ns, not significant from two-tailed unpaired t-test. (**F, G**) Immunoblot analyses of indicated proteins in COLO-800 cells. Cells were analyzed after 24 h incubation with the indicated purified proteins. Mock-purified protein was used as control. The normalized ratio between PD-L1 and TfR is shown below (**F**). **(H)** Quantification of flow cytometry analyses of cell surface PD-L1 in COLO-800 cells. n=3 biologically independent samples. Data are displayed as mean ± SD. ****P<0.0001 from two-tailed unpaired t-test. **(I)** qRT-PCR analysis of *PD-L1* expression in COLO-800 cells. Cells were analyzed after 24 h incubation with the indicated purified proteins. n=4 biologically independent samples. Data are displayed as mean ± SD. ns, not significant from two-tailed unpaired t-test. **(J)** TOPflash assay with HEK293T cells. Cells were analyzed after 12 h incubation with indicated proteins. n=3 biologically independent samples. Data are displayed as mean ± SD. ns, not significant; ****P<0.0001 from two-tailed unpaired t-test. 0.027, 0.081, 0.27, 0.81 and 2.7 nM protein used in (**A**). 1 nM protein used in (**B-E**). 0.081, 0.27, 0.81 and 2.7 nM protein used in (**F**). 0.81 nM protein used in (**G-I**). 0.12, 0.36, 1.08 and 3.24 nM RSPO2 or R2PD1 in (**J**). TfR, transferrin receptor. See also **Figure S3 and S4**.

The expression of PD-L1 is regulated by various cytokines residing in the tumor microenvironment, notably Interferon-Gamma (IFNγ) (Mandai et al., 2016; Mimura et al., 2018). We asked whether R2PD1 is capable to degrade PD-L1 induced by IFNγ. Expectedly, IFNγ increased PD-L1 protein level and co-incubation with R2PD1 decreased it (Figure S3D).

To rule out that reduction of PD-L1 was due to its transcriptional misregulation rather than induced protein degradation, we monitored PD-L1 mRNA in MEL624 cells. qRT-PCR analysis revealed no significant changes of *PD-L1* mRNA in cells incubated with the effector proteins under study (Figure 3E).

We corroborated our finding with another melanoma cell line, COLO-800, where PD-L1 is expressed and can be reduced by siRNA transfection (Figure S3E). Once again, R2PD1 decreased both the cell surface- and the total protein of PD-L1, although less complete (50-70% reduction) than in MEL624 cells (85-90% reduction) (Figures 3F-3H). Again, no significant change of *PD-L1* mRNA was detected by qRT-PCR analysis (Figure 3I). Despite the comparable protein levels of PD-L1 in MEL624 and COLO-800 cells (Figure S3G), the reduction of PD-L1 in COLO-800 was evident only after 24 hours incubation with R2PD1 chimera (Figure S3F), later than with MEL624 cells (Figure S3C). We speculated that lower efficiency and distinct kinetics of PD-L1 degradation by R2PD1 is due to the different expression levels of ZNRF3/RNF43. Indeed, qRT-PCR analysis revealed significantly higher expression of *ZNRF3/RNF43* in MEL624-than in COLO-800 cells (Figure S3H).

Atezolizumab is a therapeutic monoclonal antibody targeting PD-L1 and blocking its interaction with PD-1, thereby promoting tumor-directed T-cell activation (Herbst et al., 2020; Herbst et al., 2014). Here, we compared the effect of RSPO chimera and Atezolizumab on PD-L1 protein levels. R2PD1 chimera again reduced PD-L1 in both MEL624 and COLO-800 cells, while addition of Atezolizumab did not change PD-L1 protein levels (Figures S3I and S3J). Collectively, the data support that R2PD1 chimera degrades endogenous PD-L1 protein in melanoma cancer cells.

### PD-L1 degradation by RSPO chimera is independent of WNT/β-cat and BMP signaling

RSPOs are potent WNT signaling agonists and BMP signaling antagonists. Since these signaling activities may lead to unwanted side effects when applying an RSPO chimeric protein, the here used R2PD1 protein is deleted of its TSP1 domain, which completely abolishes its BMP inhibition (Lee et al., 2020) and it contains a F^109^A (FA) mutation within the FU2 domain, that should abrogate LGR-binding (Figure S1) and hence should prevent WNT activation. To rule out that the R2PD1 furin domains retain residual WNT/β-cat signaling activity, we performed PD-L1 degradation assays in presence of WNT inhibitors. R2PD1 chimera degraded PD-L1 both in presence of the potent WNT signaling inhibitor DKK1 (Figures S4A and S4B) and upon *β-catenin* siRNA knockdown (Figures S4C and S4D). Moreover, the chimeric protein failed to induce expression of the WNT targeted gene *AXIN2* (Figure S4E) or affect WNT reporter (TOPflash) activity, unlike wild-type RSPO2 (Figure 3J). To corroborate disabled BMP inhibition by R2PD1, we analyzed Smad1 phosphorylation, a hallmark of activated BMP signaling. Unlike wildtype RSPO2, R2PD1 failed to inhibit BMP4-mediated increase of phospho-Smad1 (Figure S4F). These findings corroborate that the RSPO chimera is WNT- and BMP signaling-disabled and that PD-L1 degradation is independent of WNT/β-cat and BMP signaling.

### PD-L1 degradation by RSPO chimera requires ZNRF3/RNF43

To address whether PD-L1 degradation via R2PD1 proceeds through ZNRF3/RNF43, we incubated R2PD1 chimeric protein with a soluble extracellular domain of RNF43 (RNF43^ECD^Fc) before adding it to cells (Figure 4A). Provided that the chimera acts specifically, RNF43^ECD^Fc should nullify the effect of R2PD1. Consistently, R2PD1 pre-incubated with RNF43^ECD^Fc rescued PD-L1 from degradation (Figure 4B). Furthermore, while single siRNA knockdown of either *ZNRF3* or *RNF43* was not sufficient to block the degradation of PD-L1 by R2PD1, double knockdown successfully prevented the degradation in both MEL624 (Figures 4C-4E) and COLO-800 cells (Figures 4F and 4G), indicating that the chimera engages both E3 ligases. To corroborate these findings, we utilized a *ZNRF3/RNF43* double-mutant A375 melanoma cell line (dKO) (Radaszkiewicz et al., 2021). While R2PD1 reduced PD-L1 in control A375 cells, it failed to decrease PD-L1 in dKO cells (Figures 4H and 4I). We conclude that in melanoma cells, RSPO chimera engages ZNRF3 and RNF43 for PD-L1 degradation.

**Figure 4.**
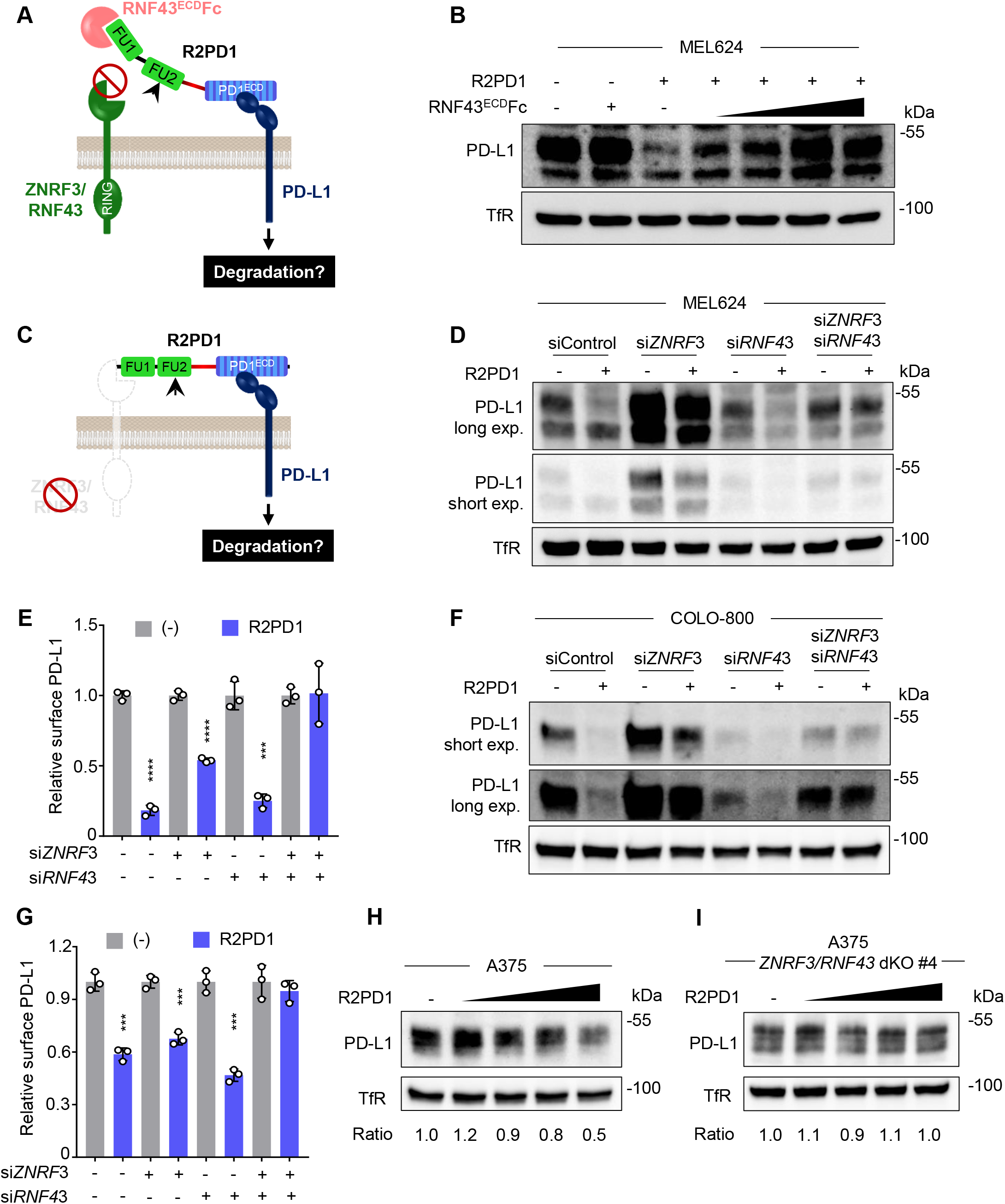
PD-L1 degradation by RSPO chimera requires ZNRF3/RNF43. **(A)** Scheme of PD-L1 degradation assay in (**B**). PD-L1 degradation by R2PD1 in presence of competing soluble RNF43^ECD^ Fc. **(B)** Immunoblot analysis of indicated proteins in MEL624 cells. Cells were analyzed after 24 h incubation with the indicated proteins. **(C)** Scheme of PD-L1 degradation assays in (**D-G**). PD-L1 degradation by R2PD1 after *ZNRF3/RNF43* knock down. **(D)** Immunoblot analysis of indicated proteins in MEL624 cells. Cells were analyzed after 24 h incubation with the indicated proteins. Cells were transfected with indicated siRNAs for 2 days before incubation with purified protein. **(E)** Quantification of flow cytometry analysis of cell surface PD-L1 in MEL624 cells. Values are normalized to the corresponding control. n=3 biologically independent samples. Data are displayed as mean ± SD. ***P<0.001, ****P<0.0001 from two-tailed unpaired t-test. **(F)** Immunoblot analysis of indicated proteins in COLO-800 cells. Cells were analyzed after 24 h incubation with the indicated proteins. Cells were transfected with indicated siRNAs for 2 days before incubation with purified protein. **(G)** Quantification of flow cytometry analysis of cell surface PD-L1 in COLO-800 cells. Values are normalized to the corresponding control. n=3 biologically independent samples. Data are displayed as mean ± SD. ***P<0.001 from two-tailed unpaired t-test. (**H, I**) Immunoblot analyses of indicated proteins in A375 cells. Cells were analyzed after 24 h incubation with the indicated proteins. 50, 150, 500 and 1,500 ng ml^-1^ RNF43^ECD^ Fc was used in (**B**). 0.27 nM (**B**), 0.24 nM (**D**), 1 nM (**E**), 0.8 nM (**F**), 2.5 nM (**G**) purified R2PD1 used. 0.073, 0.24, 0.73 and 2.44 nM (**H, I**) purified R2PD1 used.

## DISCUSSION

We introduce ROTACs, signaling-disabled R-spondin chimeras as a novel strategy for targeted degradation of transmembrane proteins. As a proof of concept, we designed and validated the R2PD1 chimeric protein to target the highly relevant immune checkpoint protein PD-L1 and demonstrate its degradation in three melanoma cell lines. Importantly, PD-L1 degradation occurred in the pico-to low nanomolar range, indicating that the chimera is highly potent. While ROTACs will be applicable only to cells that express *ZNRF3* or *RNF43*, both genes show broad to ubiquitous expression across many tissues and cell lines (https://www.ncbi.nlm.nih.gov/gene/84133 and -54894).

The R2PD1 chimera is mostly based on human proteins, reducing the risk of immunogenicity. While conceptually degradation of PD-L1 should be more efficient in overcoming immune checkpoint inhibition than antibody neutralization, a direct comparison will require *in vivo* analysis or *in vitro* tests in presence of cytotoxic T-cells. We envisage ROTACs to be complementary to LYTACs that are based on recycling receptors (Ahn et al., 2021; Banik et al., 2020; Zhou et al., 2021) or chimeric antibodies (Cotton et al., 2021; Marei et al., 2022).

The results indicate that the efficacy of the ROTACs may be limited by two main factors, the affinity for the targeted protein and the expression levels of *ZNRF3* and *RNF43* in the target cells. With regard to E3 ligase expression levels, the fact that R2PD1 degraded on the one hand even highly overexpressed PD-L1 provided that *ZNRF3/RNF43* were cotransfected, and that on the other hand endogenous PD-L1 degradation was more efficient in MEL624-than in COLO-800 that express high and low *ZNRF3/RNF43* levels, respectively, suggests that target cancer cells with high *ZNRF3/RNF43* expression may be preferable for R2PD1 chimera application. Conversely, only dual knockdown of both *ZNRF3* and *RNF43* completely abolished PD-L1 degradation in MEL624 and COLO-800 cells, supporting that both E3 ligases act redundantly. We envision that the modular nature of the here introduced WNT- and BMP signaling incompetent ROTACs will find applications beyond PD-L1.

## ACKNOWLEDGEMENTS

We thank Vita Bryja and Rienk Offringa for reagents, Andrey Glinka for critical discussion. Expert technical support by the DKFZ core facilities for light microscopy and flow cytometry is gratefully acknowledged. This work was supported by German Research Council (DFG) via the Collaborative Research Centre 1324 TP B1.

## AUTHOR CONTRIBUTIONS

R.S. and C.N. conceived the project. R.S. designed and performed the experiments. H.L. performed part of experiments and provided cartoons. R.S. analyzed the data with input from C.N.. R.S. and C.N. wrote the manuscript.

## METHODS

### Resource availability

#### Lead Contact and Materials Availability

Further information and requests for materials are directed to the Lead Contact, Christof Niehrs.

#### Materials Availability Statement

All unique materials newly generated in this study are available from the lead contact with a completed Materials Transfer Agreement.

### Experimental Model and Subject Details

#### Cell lines and growth conditions

HEK293T, MEL624 and A375 cells were maintained in DMEM High glucose (Gibco 11960) supplemented with 10% FBS (Capricorn FBS-12A), 1% penicillin-streptomycin (Sigma P0781), and 2mM L-glutamine (Sigma G7513). A375 wild type and *ZNRF3/RNF43* double knockout cells were gifts from Vita Bryja (Radaszkiewicz et al., 2021). COLO-800 cells were maintained in RPMI (Gibco 21875) with 10% FBS, 1% penicillin-streptomycin, 2mM L-glutamine and 1mM sodium pyruvate (Sigma S8636). All cell lines were cultured at 37°C and 5% CO2 in a humidity-controlled incubator. Mycoplasma contamination was negative in all cell lines used.

### Method details

#### Constructs

Human RSPO2 wild type construct C-terminally tagged with Flag in pCS2+ vector was previously reported (Lee et al., 2020; Sun et al., 2021). R2^FA^ with the phenylalanine-alanine mutation at position of 109 within FU2 domain was obtained by mutagenesis PCR. PD1 was cloned by inserting the N-terminus to the end of the IgV-like domain of human PD-1 in pCS2+ vector. The high-affinity consensus mutant of PD-1 (PD1^HAC^) (Maute et al., 2015) was generated by Gibson assembly with synthetic oligoes. Human PD-L1 extracellular domain was inserted into the alkaline phosphatase (AP)-pCS2+ vector and expressed as the AP fusion protein (PD-L1^ECD^AP). RSPO chimeras R2PD1 and R2^wt^PD1 were obtained through Gibson assembly with fragments from the FU1/2 domains of R2^FA^/RSPO2 and PD1^HAC^ constructs and inserted into pCS2+ and AP-pCS2+ vectors. A 10-amino acid glycine-serine linker (GSGSGGSGSG) was inserted between RSPO2 FU2 domain and PD1^HAC^. Human ZNRF3-HA, ZNRF3^ΔRING^-myc and RNF43-Flag constructs were described previously (Chang et al., 2020; Kim et al., 2021). Human PD-L1-GFP (pEGFP-N1/PD-L1) was a gift from Mien-Chie Hung (Addgene plasmid # 121478; http://n2t.net/addgene:121478; RRID: Addgene_121478) (Li et al., 2016). The sequences of the constructs generated in this study were confirmed by DNA sequencing.

#### Cell transfection

siRNAs and plasmids were transfected using DharmaFECT 1 transfection reagent (Dharmacon T-2001) and X-tremeGENE9 DNA transfection reagent (Roche 06365809001) respectively, according to the manufacturer.

#### Generation of conditioned medium and protein purification

HEK293T cells were seeded into 10 cm culture dishes and transiently transfected with pCS2+ empty vector, RSPO2-Flag, R2^FA^-Flag, PD1^ECD^-Flag, PD1^HAC^-Flag, R2PD1-Flag, PD-L1^ECD^AP, R2PD1-AP and R2^wt^PD1-AP. After 24 hours, media were changed to fresh DMEM containing 10% FBS, 1% L-glutamine and 1% penicillin-streptomycin, and harvested daily in the following three days. Conditioned media were validated and quantified by immunoblot. Media containing equal mole amount of proteins were used in the subsequent analysis. WNT3A conditioned medium was produced in L-cells as previously described^11^.

For protein purification, conditioned media were over-night incubated with anti-Flag antibody conjugated agarose beads (Sigma A2220). After washing with ice-cold PBS, proteins bound to the beads were eluted by 200 mM Glycine (pH 2.6) and neutralized with equal volume of 1 M Tris (pH 8.0). The eluate was dialyzed against ice-cold PBS with at least 1000-fold volume excess. Protein concentration was quantified by Coomassie staining using BSA standards (Sigma P0914).

#### *In vitro* binding assay

High binding 96-well plates (Greiner M5811) were coated with 2 μg ml^-1^ of recombinant human PD-L1^ECD^Fc (Peprotech 310-35), ZNRF3^ECD^Fc (R&D systems 7994-RF-025) or RNF43^ECD^Fc (R&D systems 7964-RN-050) protein in bicarbonate coating buffer (50 mM NaHCO3, pH 9.6) overnight at 4 °C. Coated wells were washed three times with TBST (TBS, 0.1% Tween-20) and blocked with 5% BSA in TBST for 1 hour at room temperature. After overnight incubation with Flag tagged proteins, wells were washed three times with TBST. Protein bound to the wells were detected with a peroxidase conjugated anti-Flag antibody (Sigma A8592) and quantified with QuantaBlu™ Fluorogenic Peroxidase Substrate Kit (Thermo Scientific™ 15169). For the ternary complex formation assays, instead of HRP-anti-Flag antibody, PD-L1^ECD^AP was added to wells. After overnight incubation, AP signal was detected with the chemiluminescent AquaSpark AP substrate (Serva 42593.01). Data are displayed as average of biological replicates with SD.

#### Cell surface binding assay

Cell surface–binding assays were done as previously described^20^. In brief, HEK293T cells were transfected with *LGR4* DNA and incubated with conditioned media for 3 hours. Surface binding was detected by development with BM-Purple (Sigma 11442074001). Images were obtained with DMIL microscope/Canon DS126311 camera (LEICA) and processed with ImageJ.

#### PD-L1 degradation assay

HEK293T cells were seeded into cell culture plates and transfected with indicated plasmids. 24 hours after transfection, cells were treated with either conditioned media or purified proteins as indicated. After another 24 hours incubation, cells were harvested for immunoblot.

Cancer cells were seeded into cell culture plates and treated as indicated. After 24 hours incubation or indicated periods, cells were harvested for immunoblot. For the degradation assay in present of IFNγ stimulation, MEL624 cells were pre-incubated with 333 U IFNγ (ImmunoTools 11343536) for 24 hours. PD-L1 antibody Atezolizumab and matched isotype control used in the assay were gifts from Offringa lab from DKFZ, Germany.

#### Immunoblot

Cultured cells were harvested and lysed in ice-cold RIPA buffer with cOmplete Protease Inhibitor Cocktail (Roche 11697498001). Lysates were mixed with Laemmli buffer containing β-mercaptoethanol and boiled at 70 °C for 10 min to prepare SDS-PAGE samples. For cytosolic β-catenin detection, saponin buffer was used (Acebron et al., 2014). Immunoblot images were acquired with SuperSignal West pico ECL (ThermoFisher 34580) using LAS-3000 system (FujiFilm). Quantification of blots was done using ImageJ software.

#### Immunofluorescence (IF)

Cells were seeded to cell culture plates with inserted glass coverslips. 24 hours after transfection with indicated plasmids, cells were treated with purified proteins for indicated periods. After fixation with 4% PFA for 10 min, cells were stained with primary antibodies (1:250) overnight at 4 °C, and fluorophore conjugated secondary antibodies (1:250) at room temperature for 2 hours. Images were obtained using LSM 700 (Zeiss) and analyzed with ImageJ. Around 100 cells were analyzed in each group.

#### Flow cytometry

Cells were harvested through a none-enzymatic approach, pelleted and resuspended in ice-cold blocking buffer (PBS supplemented with 1%BSA and 0.1% NaN_3_). After blocking with Fc receptor binding inhibitor (eBioscience 14916173), cells were staining with anti-PD-L1 antibody (CST 86744), followed by incubation with a fluorochrome-labled secondary antibody. Staining with secondary antibody only was used as control. Dead cells were excluded by counterstaining with propidium iodide. FACS Samples were analyzed with FACSCanto or LSRFortessa (BD Biosciences) and data were processed with FlowJo software. Rabbit anti-PD-L1 (CST 86744) antibody was used for cell surface PD-L1 measurement.

#### Quantitative real-time PCR

Cultured cells were lysed in Macherey-Nagel RA1 buffer containing 1% β-mercaptoethanol and total RNAs were isolated using NucleoSpin RNA isolation kit (Macherey-Nagel 740955). Reverse transcription and PCR amplification were performed as described before^21^. Primers used for *AXIN2* were: forward 5’-CCACACCCTTCTCCAATCC-3’ and reverse 5’-TGCCAGTTTCTTTGGCTCTT-3’. Primers used for *PD-L1* were: forward 5’-CCTACTGGCATTTGCTGAACG-3’ and reverse 5’-AGACAATTAGTGCAGCCAGGT-3’. Primers used for *ZNRF3* were: forward 5’-TGTGCCATCTGTCTGGAGAA-3’ and reverse 5’-TTCCTGTGAAACCGGTGAGT-3’. Primers used for *RNF43* were: forward 5’-GTTTGCTGGTGTTGCTGAAA-3’ and reverse 5’-TGGCATTGCACAGGTACAG-3’. Primers used for *GAPDH* were: forward 5’-AGCCACATCGCTCAGACAC-3’ and reverse 5’-GCCCAATACGACCAAATCC-3’. Graphs show relative gene expressions to *GAPDH*. Data are displayed as mean with SD from multiple replicates.

#### TOPflash luciferase reporter assay

TOPflash luciferase assays were carried out as previously described (Berger et al., 2017). Data are displayed as average of biological replicates with SD.

#### Quantification and statistical analysis

Statistical analyses were done with the PRISM7 software using unpaired t-test or one-way ANOVA test. Not significant (ns), p > 0.05, *p < 0.05, **p < 0.01, ***p < 0.001, ****p < 0.0001.

## SUPPLEMENTAL FIGURE TITLES AND LEGENDS

**Figure S1.**
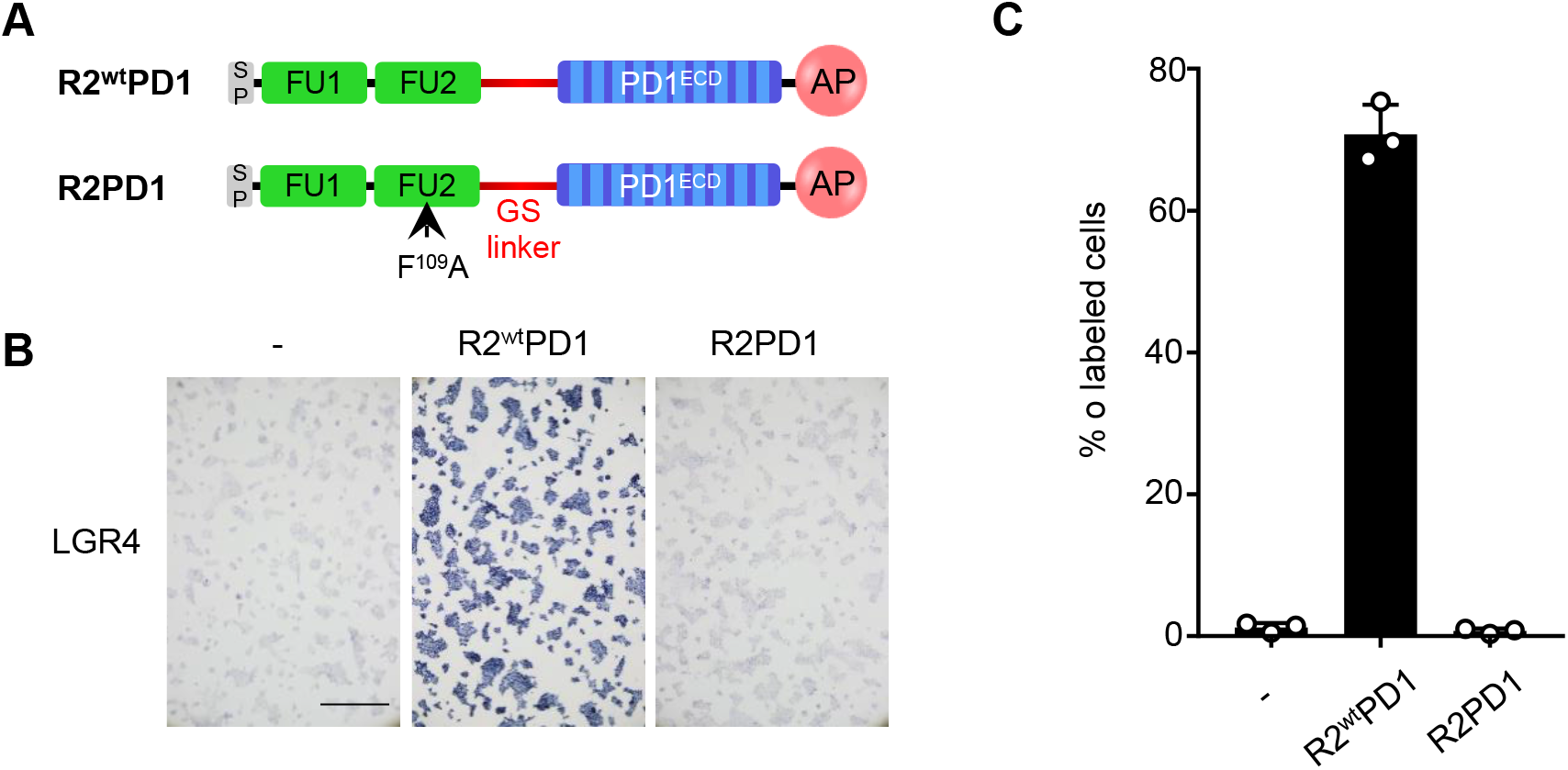
RSPO chimera with F^109^A mutation does not bind LGR4. (**A)** Domain-structure of indicated proteins. SP, signal peptide; FU, furin domain; GS, glycine-serine linker; ECD, extracellular domain; wt, wild-type; AP, alkaline phosphatase. (**B**-**C**) Cell surface binding assay in HEK293T cells. Cells were transfected with *LGR4* and treated with same amount of indicated AP tagged proteins. Binding was detected as purple stain on cell surface by chromogenic AP assay. Quantification shown in (**C**). Scale bar, 0.5 mm. n=3 experimentally independent samples. Data are displayed as mean ± SD.

**Figure S2.**
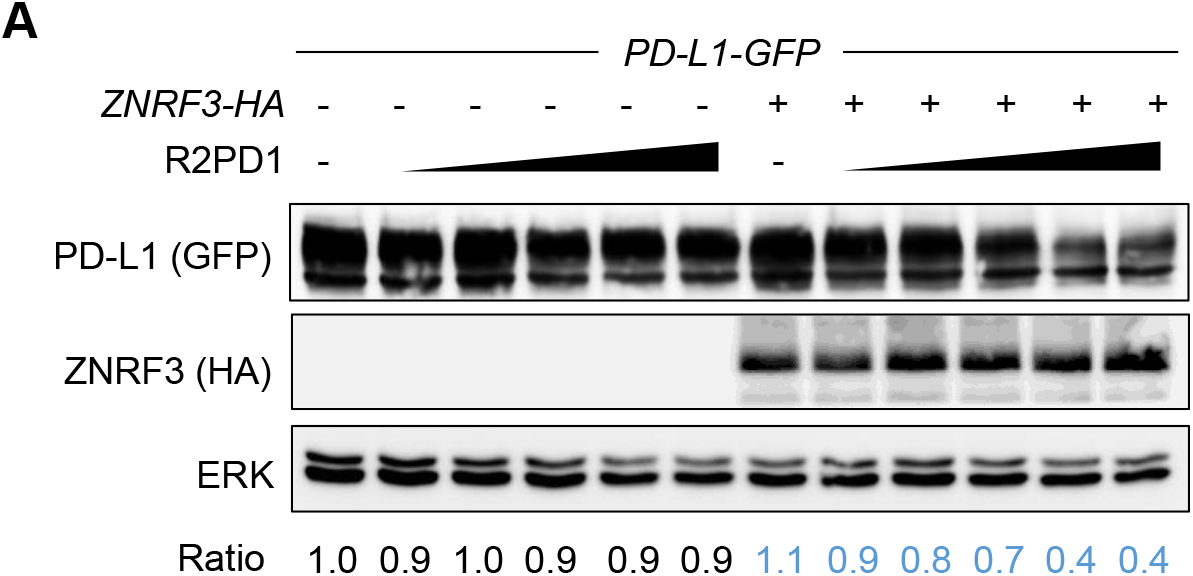
R2PD1 promotes degradation of overexpressed PD-L1. (**A**) Immunoblot analysis of indicated proteins in HEK293T cells. Cells were transfected for 2 days with *PD-L1-GFP* with or without *ZNRF3-HA*. Cells were analyzed after 24 h incubation with purified R2PD1 protein (0.2, 0.8, 2.2, 5.4 and 13.6 nM). The normalized ratio between PD-L1 and ERK is shown below.

**Figure S3.**
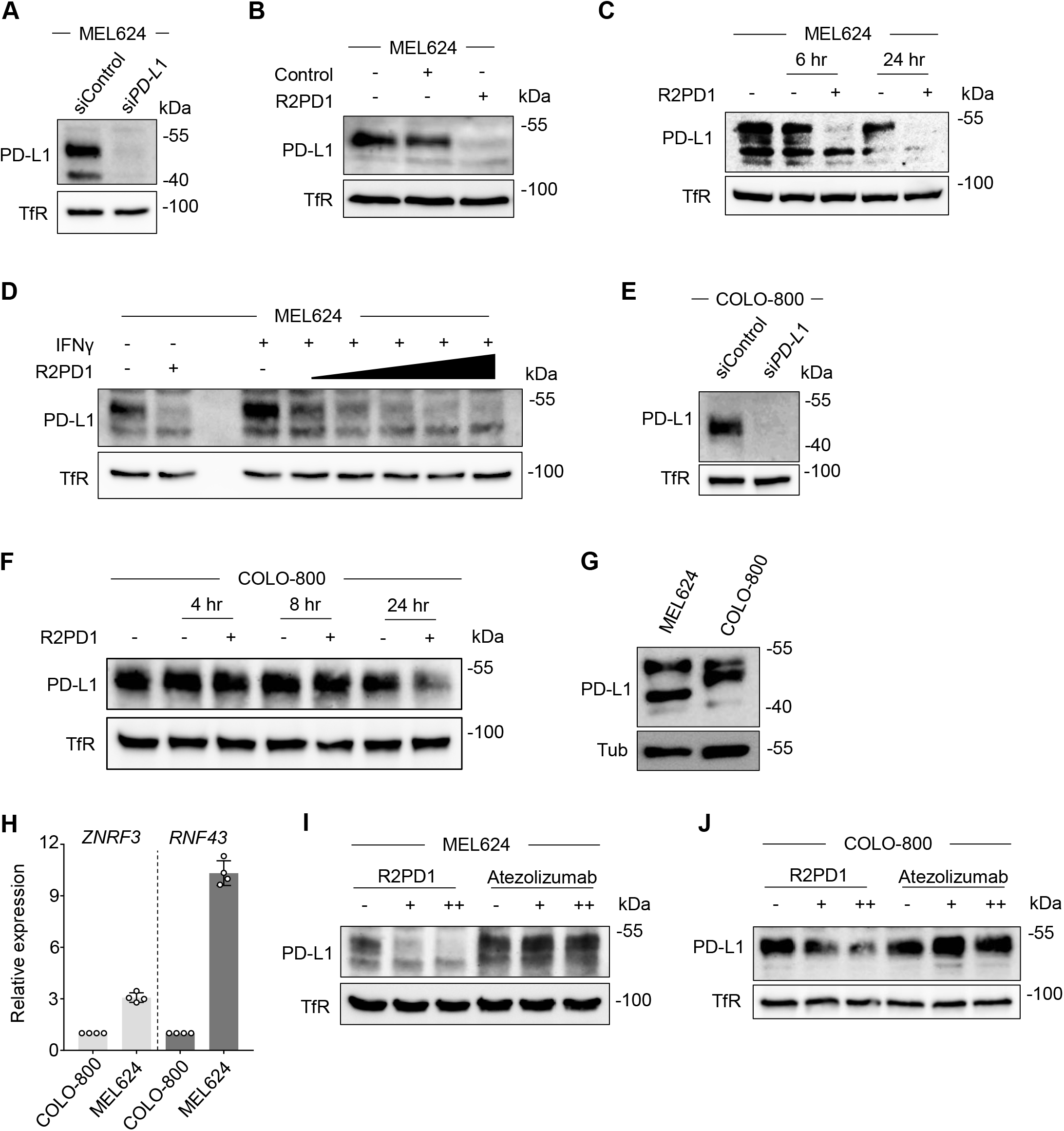
RSPO chimera promotes degradation of endogenous PD-L1 in melanoma cells. (**A-G**) Immunoblot analyses of indicated proteins in MEL624 and COLO800 cells as indicated. **(A)** Cells were transfected for 3 days with indicated siRNAs. **(B)** Cells were analyzed after 24 h incubation with indicated conditioned medium. **(C)** Cells were treated with 0.24 nM purified proteins for the indicated hours before analysis. **(D)** Cells were treated with 333 U IFN-γ for 24 h and harvested for analysis after additional 24 h incubation with purified R2PD1 proteins (0.027, 0.081, 0.27, 0.81 and 2.7 nM). **(E)** Cells were transfected for 3 days with indicated siRNA before analysis. **(F)** Cells were analyzed after incubation with 0.7 nM purified R2PD1 for the indicated hours. **(G)** Untreated cells were analyzed. **(H)** qRT-PCR analysis of *ZNRF3/RNF43* expression in MEL624 and COLO-800 cells. n=4 experimentally independent samples. Data are displayed as mean ± SD. (**I, J**) Immunoblot analyses of indicated proteins in MEL624 and COLO800 cells as indicated. Cells were analyzed after 24 h incubation with purified R2PD1 or anti-PD-L1 antibody (Atezolizumab). (**I**) 0.07 and 0.24 nM, (**J**) 0.07 and 0.7 nM. Mock-purified protein and isotypic antibody control were used as controls.

**Figure S4.**
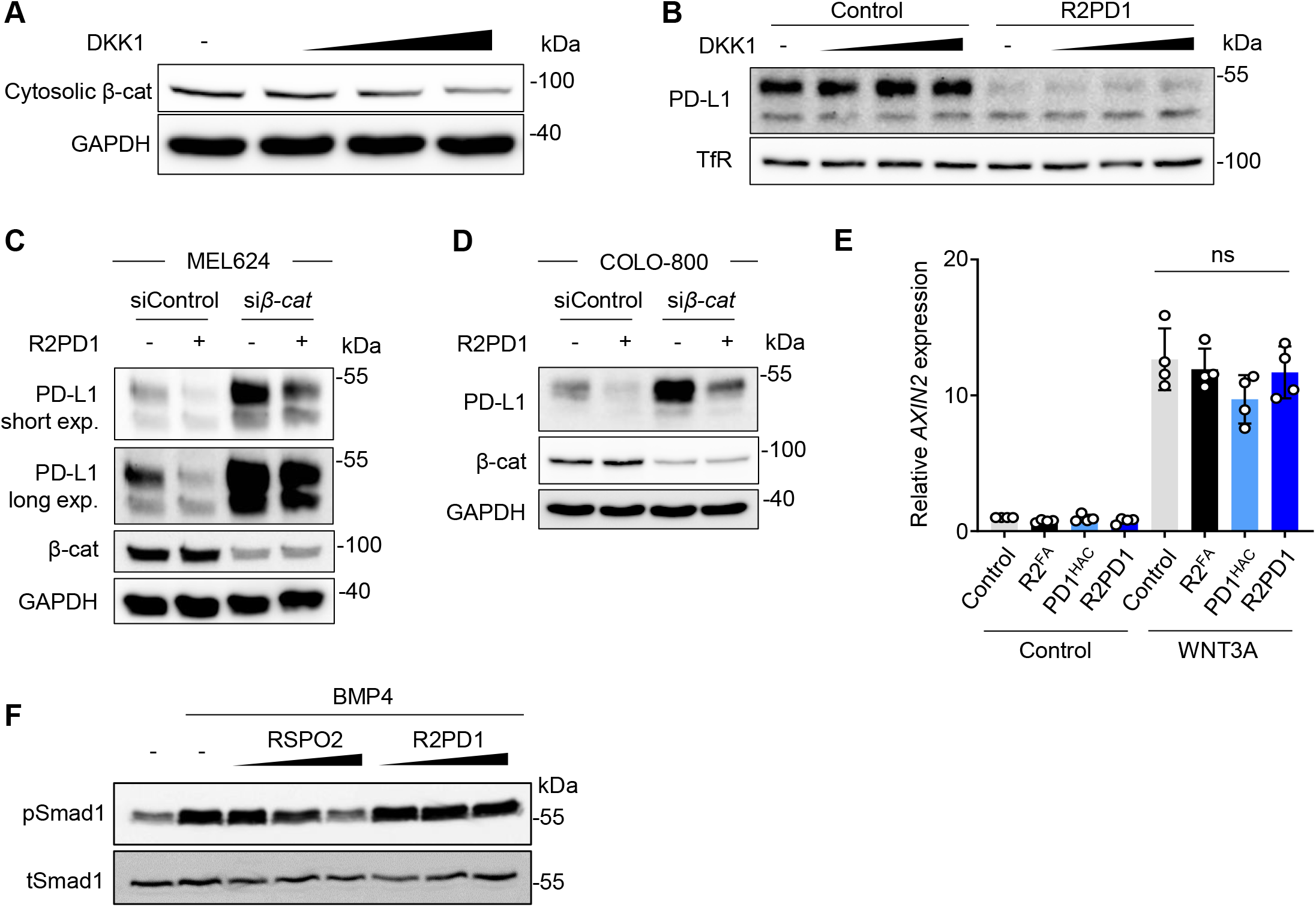
PD-L1 degradation by RSPO chimera is independent of WNT/β-cat and BMP signaling. (**A-D**) Immunoblot analyses of indicated proteins in MEL624 (**A-C**) and COLO-800 (**D**) cells. Cells were analyzed after 12 h (**A**) or 24 h (**B**-**D**) incubation with indicated proteins. Cells were transfected with indicated siRNAs for 2 days before incubation with purified protein (**C**-**D**). **(E)** qRT-PCR analysis of *AXIN2* expression in COLO-800 cells. Cells were analyzed after 24 h incubation with purified proteins in presence of WNT3A. n=4 biologically independent samples. Data are displayed as mean ± SD. ns, not significant from two-tailed unpaired t-test. **(F)** Immunoblot analysis of indicated proteins in MEL624 cells. Cells were analyzed after 1 h incubation with indicated proteins. 10, 50 and 200 ng ml^-1^ DKK1 protein used in (**A, B**). Purified R2PD1 used with 0.24 nM (**B, C**), 0.73 nM (**D**), 0.8 nM (**E**).

